# Stromal CTHRC1 protects the valvular interstitium from macrophage-associated inflammatory remodeling and calcification

**DOI:** 10.64898/2026.09.21.753362

**Authors:** Hiroshi Sakamoto, Tomohisa Sakaue, Mika Hamaguchi, Yasuhisa Nakao, Mie Kurata, Akiyasu Iwase, Hiroki Kurihara, Yosuke Kiriyama, Takuma Fukunishi, Hirotsugu Kurobe, Takashi Nishimura, Shunji Uchita, Daisuke Shiokawa, Yoshiaki Kubota, Osamu Yamaguchi, Hironori Izutani

**Affiliations:** Department of Cardiovascular and Thoracic Surgery, Graduate School of Medicine, Ehime University, Shitsukawa, Toon City, Ehime, 791-0295, Japan; Department of Tumor Microenvironment Regulation, Next-Generation Precision Cancer Research Center, Osaka International Cancer Institute, 3-1-69 Otemae, Chuo-ku, Osaka 541-8567, Japan; Department of Cardiology, Pulmonology, Nephrology and Hypertension, Graduate School of Medicine, Ehime University, Shitsukawa, Toon City, Ehime, 791-0295, Japan; Department of Analytical Pathology, Graduate School of Medicine, Ehime University, Shitsukawa, Toon City, Ehime, 791-0295, Japan; Isotope Science Center, The University of Tokyo, 2-11-16 Yayoi, Bunkyo-ku, Tokyo 113-0032, Japan; Laboratory of Multicellular Dynamics, International Research Center for Medical Sciences, Kumamoto University, 2-2-1 Honjo, Chuo-ku, Kumamoto 860-0811, Japan; Translational Research Center, Ehime University Hospital, Shitsukawa, Toon City, Ehime, 791-0295, Japan; Department of Anatomy, Keio University School of Medicine, 35 Shinanomachi, Shinjuku-ku, Tokyo 160-8582, Japan

**Keywords:** Calcific aortic valve disease, Valvular interstitial cells, CTHRC1, Valvular calcification

## Abstract

**Background:** Calcific aortic valve disease (CAVD) is characterized by progressive inflammatory and fibrocalcific remodeling. Although valvular interstitial cells (VICs) are generally considered to drive fibrosis and osteogenic remodeling, whether injury-activated VICs mount endogenous protective responses that preserve the valvular interstitial microenvironment and restrain calcification remains unknown.

**Methods:** We performed spatial transcriptomic profiling of aortic valves in a mouse model of endothelial injury-induced CAVD to define early injury-responsive programs within the valvular interstitium. The cellular origin and spatial distribution of candidate protective factors were examined by immunohistochemistry and lineage tracing, and their relevance to human disease was assessed using stenotic aortic valves. The functional role of CTHRC1 was investigated using genetic *Cthrc1* deficiency combined with longitudinal hemodynamic assessment, histological analysis, and spatial transcriptomic profiling.

**Results:** Spatial transcriptomics identified *Cthrc1* as a prominent component of an early injury-induced stromal response in the expanding valvular interstitium. CTHRC1 was strongly expressed in activated VICs within thickened murine valve leaflets and human stenotic aortic valves. Lineage tracing demonstrated that the expanded VIC population arose predominantly from PDGFRβ^+^ resident interstitial cells, with minimal endothelial contribution. Despite comparable early hemodynamic responses to endothelial injury, *Cthrc1* deficiency exacerbated chronic valvular calcification. Spatial profiling of *Cthrc1*-deficient valves revealed pronounced interstitial accumulation of galectin-3^+^ foamy macrophages, accompanied by mitochondrial respiratory-chain signature loss and cell death-associated pathway activation. These findings indicate that transient CTHRC1 induction after endothelial injury defines an endogenous stromal protective response that preserves the valvular interstitial microenvironment and limits macrophage-associated tissue injury and subsequent dystrophic calcification.

**Conclusions:** Injury-activated VICs are not merely effectors of pathological remodeling, but can engage an endogenous tissue-protective response through CTHRC1. These findings identify a previously unrecognized stromal defense mechanism linking endothelial injury to macrophage-associated inflammatory remodeling and dystrophic calcification and suggest CTHRC1-dependent stromal protection as a potential therapeutic axis for limiting CAVD progression.

## Introduction

Aortic stenosis (AS) is the most prevalent heart valve disorder in developed countries and predominantly affects older adults. In a large European prospective survey of patients with valvular heart disease, AS was the most frequent native valve lesion (1). In AS, progressive leaflet thickening reduces cusp pliability and narrows the valve orifice, imposing a chronic pressure overload on the left ventricle, which culminates in decompensated heart failure. The severest and end-stage manifestation of AS is calcific aortic valve disease (CAVD) (1). In the PARTNER 1B trial, patients with severe, inoperable AS assigned to standard treatment had a 5-year all-cause mortality rate of 93.6%, indicating an extremely poor long-term prognosis without definitive aortic valve intervention (2).

However, no effective disease-modifying therapy exists; management ultimately relies on invasive and costly interventions, such as surgical aortic valve replacement (SAVR) or transcatheter aortic valve implantation (TAVI) (3,4). One of the challenges hindering the practical development of prophylactic and therapeutic drugs is an insufficient understanding of the molecular mechanisms underlying valve leaflet thickening and calcification in early-stage AS/CAVD.

Human aortic valve specimens collected during SAVR offer a critical window into the pathology of CAVD. In particular, single-cell RNA sequencing (scRNA-seq) of end-stage lesions has revealed profound cellular and transcriptional heterogeneity, highlighting the dynamic molecular programs that drive disease progression (5,6). scRNA-seq of human CAVD valves revealed pronounced cell-state heterogeneity with diverse immune clusters, and pseudotime analyses of valvular endothelial cells (VECs) suggested early fibrosa-biased endothelial reprogramming with endothelial-to-mesenchymal transition (EndMT)-associated signatures inferred *in silico* (5,6). While these *ex vivo* approaches have yielded important insights, they primarily reflect the terminal stages of the disease and do not capture the dynamic processes that occur during the early or mild phases *in vivo*.

Several animal models have been developed and are widely used to overcome this limitation. Multiple studies have used *Apoe^−/−^*or *Ldlr^−/−^* mice fed a high-cholesterol diet to model dyslipidemia-driven AS/CAVD (7–10). Although dyslipidemia contributes to CAVD, the disease is multifactorial, and lipid-lowering has not been shown to attenuate valve-related outcomes in randomized trials; therefore, dyslipidemia-driven models are best interpreted as modeling one clinically relevant, but not universal, disease trajectory (11,12). The prevailing view is that oscillatory shear stress-induced endothelial injury to aortic valve leaflets allows erythrocytes, lipids, and immune cells to penetrate the valve interstitium, which in turn initiates oxidative stress and inflammation, followed by myofibroblast activation, interstitial fibrosis, and ultimately calcification (13). Honda *et al.* reported an endothelial injury-induced murine AS model that was independent of dyslipidemia (14). In this pathological model, wild-type mice fed a normal chow diet developed CAVD and successfully captured the key histopathological features of end-stage human disease. By carefully optimizing the timing of euthanasia, the wire injury (WI) method has the potential to recapitulate the earliest pathological stage of AS/CAVD *in vivo*. When appropriately selected and phenotyped, animal models can recapitulate the key features of AS/CAVD progression and allow lesion development to be studied in a more integrated physiological context (15,16).

Therapeutic intervention is likely to be most effective before overt calcification develops. Elucidating the early pathological processes that precede valvular calcification is essential for the development of disease-modifying therapies. In particular, identifying genes that are induced during the early phase following valvular injury and subsequently govern calcification progression may provide novel therapeutic targets. We employed a wire-induced valvular endothelial injury model that recapitulates early valve thickening and remodeling after endothelial damage. Furthermore, to capture molecular alterations within the valvular interstitium at a spatial resolution, we generated a spatial transcriptomic atlas of early-stage AS lesions to identify candidate master regulators involved in disease progression.

## Results

### Aortic stenosis develops within 2 weeks after valvular endothelial injury

To generate a murine model of AS for spatial transcriptomic profiling, 8-week-old wild-type C57BL/6J mice fed a normal chow diet were subjected to WI (14) (Fig. 1a). To assess the temporal onset and progression of AS and characterize early tissue remodeling during disease initiation, transthoracic echocardiography (TTE) was performed serially following the WI procedures. Color Doppler imaging in the WI group revealed turbulent blood flow in the sinus of Valsalva and ascending aorta, suggesting narrowing of the effective aortic valve orifice area (Fig. 1b).

**Fig. 1.**
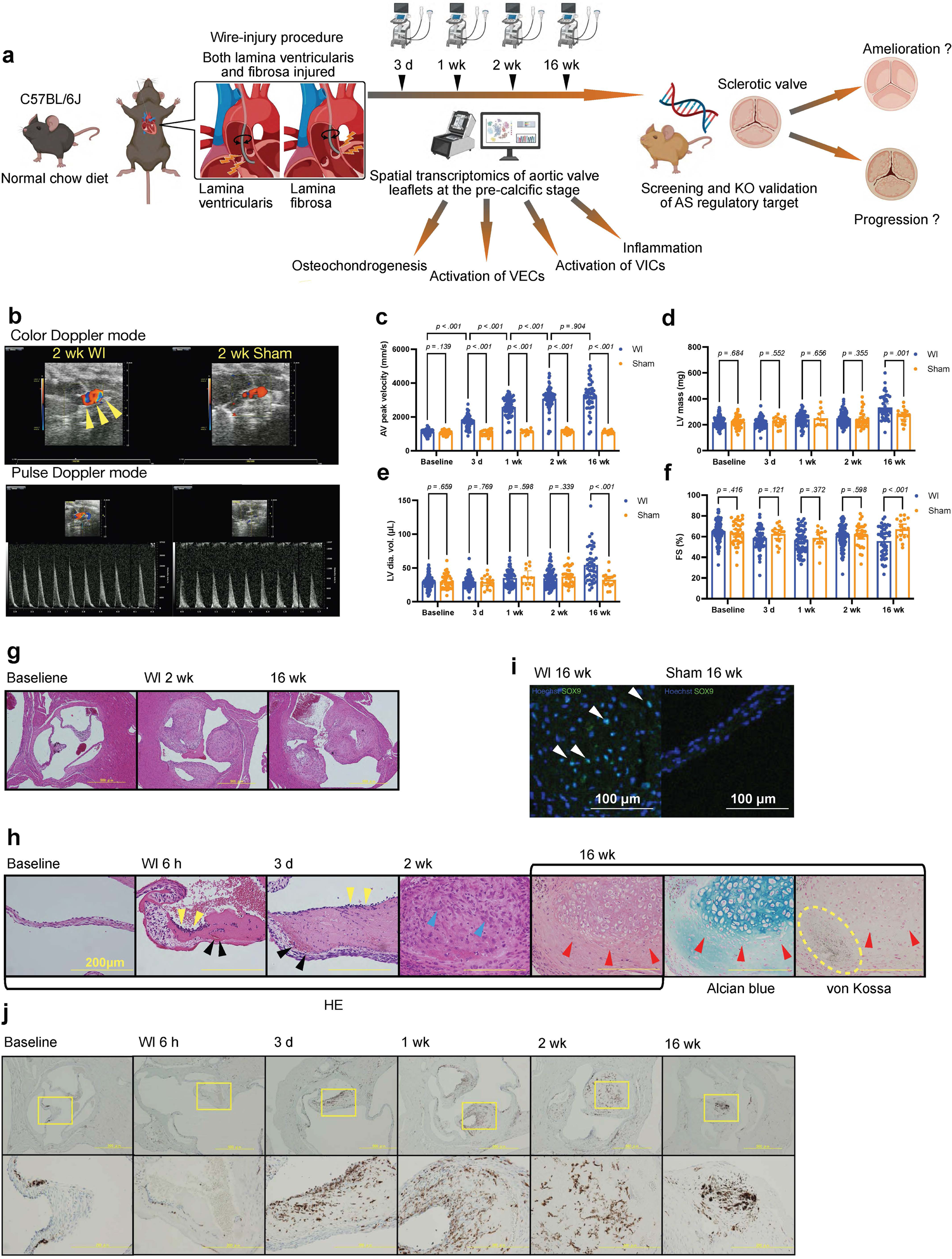
Acute-phase hemodynamic and histopathological characterization of a modified murine AS model induced by valvular endothelial injury. **a**, Schematic diagram of the study design. **b**, Representative TTE console display: top panels, color Doppler; bottom panels, pulsed-wave Doppler used to measure the peak transvalvular aortic flow velocity (AV peak velocity). Yellow arrowheads mark a mosaic aliasing jet in the sinus of Valsalva and proximal ascending aorta, consistent with a high-velocity transvalvular flow. c–f, Serial TTE in wire injury (WI) and sham mice at baseline, 3 days (d), 1 week (wk), 2 wk, and 16 wk. **c**, AV peak velocity; **d**, left ventricular (LV) mass; **e**, LV end-diastolic volume (EDV); **f**, LV fractional shortening (FS). Sample sizes per time point (n, WI/Sham): baseline 133/36; 3 d 62/17; 1 wk 61/13; 2 wk 91/32; 16 wk 45/17. The same cohort contributed to panels **c–f**; imputation was not performed for missing observations owing to scheduled euthanasia or welfare-based omissions at 3 d and 1 wk. Points denote individual mice, and bars show SD. Group × time effects were analyzed using two-way mixed-effects models (REML; sphericity not assumed; Greenhouse–Geisser correction, where applicable) with Sidak-adjusted post-hoc comparisons. Exact statistics (*F*, df, *P*, *ε*) and time-specific contrasts are provided in Supplementary Fig. 1a**–d** and Supplementary Tables 1–12. The units are indicated on the axis. g, Low-power H&E images of valves at baseline and 2 and 16 wk after WI. Scale bar, 500 µm. **h**, High-power H&E images of aortic valve leaflets at baseline, 3 d, 1 wk, 2 wk, and 16 wk after WI and Alcian blue and von Kossa staining at 16 wk. Black, yellow, and red arrowheads indicate red blood cells, mononuclear inflammatory infiltrates, and chondrocyte-like cells, respectively. Scale bar, 200 µm. **i**, High-power immunofluorescence images of SOX9 at 16 wk. Scale bar, 200 µm. **j**, Low-power (upper panels) and high-power (lower panels) immunohistochemistry images for CD68 at baseline and 3 d, 1 wk, 2 wk, and 16 wk after the WI procedure.

Longitudinal echocardiographic parameters, including aortic valve peak velocity (AV peak velocity), fractional shortening (FS), left ventricular (LV) mass, and LV end-diastolic volume (LV volume), were analyzed using linear mixed-effects models with fixed effects for group, time, and their interaction, and a random intercept for each mouse. Greenhouse–Geisser correction was applied where appropriate. AV peak velocity progressively increased after WI but remained stable in the sham group, indicating a strong time × group interaction (*F*(4, 327) = 105.3, *p* < 0.001) (Fig. 1c). The time and group main effects were also significant (*F*(2.627, 214.8) = 119.3, *p* < 0.001; *F*(1, 169) = 393.0, *p* < 0.001). These data confirmed that the WI procedure recapitulated the hemodynamic hallmarks of human AS, with peak AV velocity reaching a plateau within 2 weeks after the injury (Fig. 1c). The Sidak-adjusted time-specific contrasts are shown in Supplementary Fig. 1a and Supplementary Tables 1–3. LV mass increased over time with a significant time × group interaction (*p* = 0.002), and WI showed higher mass at 16 weeks versus earlier time points and versus sham (all Sidak-adjusted *p* < 0.05), indicating afterload-driven hypertrophy (Fig. 1d, Supplementary Fig. 1b, and Supplementary Tables 4–6). LV volume increased in the WI group, with the greatest difference observed at 16 weeks (Fig. 1e, Supplementary Fig. 1c, and Supplementary Tables 7–9). In addition, the FS declined in the WI group, reflecting afterload-related systolic impairment (Fig. 1f, Supplementary Fig. 1d, and Supplementary Tables 10–12).

In addition to the hemodynamic analysis, murine aortic valve specimens from both the WI and sham groups were harvested serially and pathologically evaluated. In low-power H&E staining, the wire-injured valves exhibited comparable leaflet thickening at 2 and 16 weeks (Fig. 1g). Notably, at 6 h and 3 days, extravasated red blood cells (RBCs), fibrin deposition (Fig. 1h, black arrowheads), and CD68^+^ cell infiltration (Fig. 1j) were observed within the valvular interstitium. Immunofluorescence (IF) analysis revealed a prominent accumulation of SOX9^+^ cells in stenotic valves at the late stage (16 weeks) following WI, indicating that the WI model recapitulates chondro-osteogenic valvular remodeling (Fig. 1i, white arrowheads). After 2 weeks, spindle-shaped fibroblast-like cells accumulated, accompanied by progressive leaflet thickening (Fig. 1h, blue arrowheads). Sixteen weeks after WI, Alcian blue staining demonstrated proteoglycan-rich chondrogenic remodeling within the valve leaflets, whereas von Kossa staining revealed prominent interstitial calcification (Fig. 1h, red arrowheads), collectively representing the key features of end-stage human AS pathology.

### Injured aortic valve leaflets exhibit osteochondrogenic characteristics

To identify early transcriptional alterations potentially underlying the subsequent calcific remodeling observed at 16 weeks, we performed spatial transcriptomic profiling of aortic valve leaflets harvested 2 weeks after WI using the 10x Genomics Visium CytAssist platform. Following quality control, six libraries were retained and integrated for downstream analysis (Supplementary Fig. 2a and b). Joint uniform manifold approximation and projection (UMAP) of spot-level transcriptomes resolved 15 clusters (Fig. 2a and Supplementary Fig. 3). Based on spatial feature mapping and H&E images, we annotated the anatomical compartments: thickened WI leaflets mapped predominantly to cluster 8, morphologically intact leaflets (including preserved regions from the WI group) to cluster 4, and the aortic annulus to cluster 10, whereas the remaining clusters represented adjacent non-valvular tissues (Supplementary Fig. 3). Consistent with the histology at this precalcific stage, we focused subsequent analyses on comparisons between clusters 8 and 4, representing early disease and reference tissue, respectively.

**Fig. 2.**
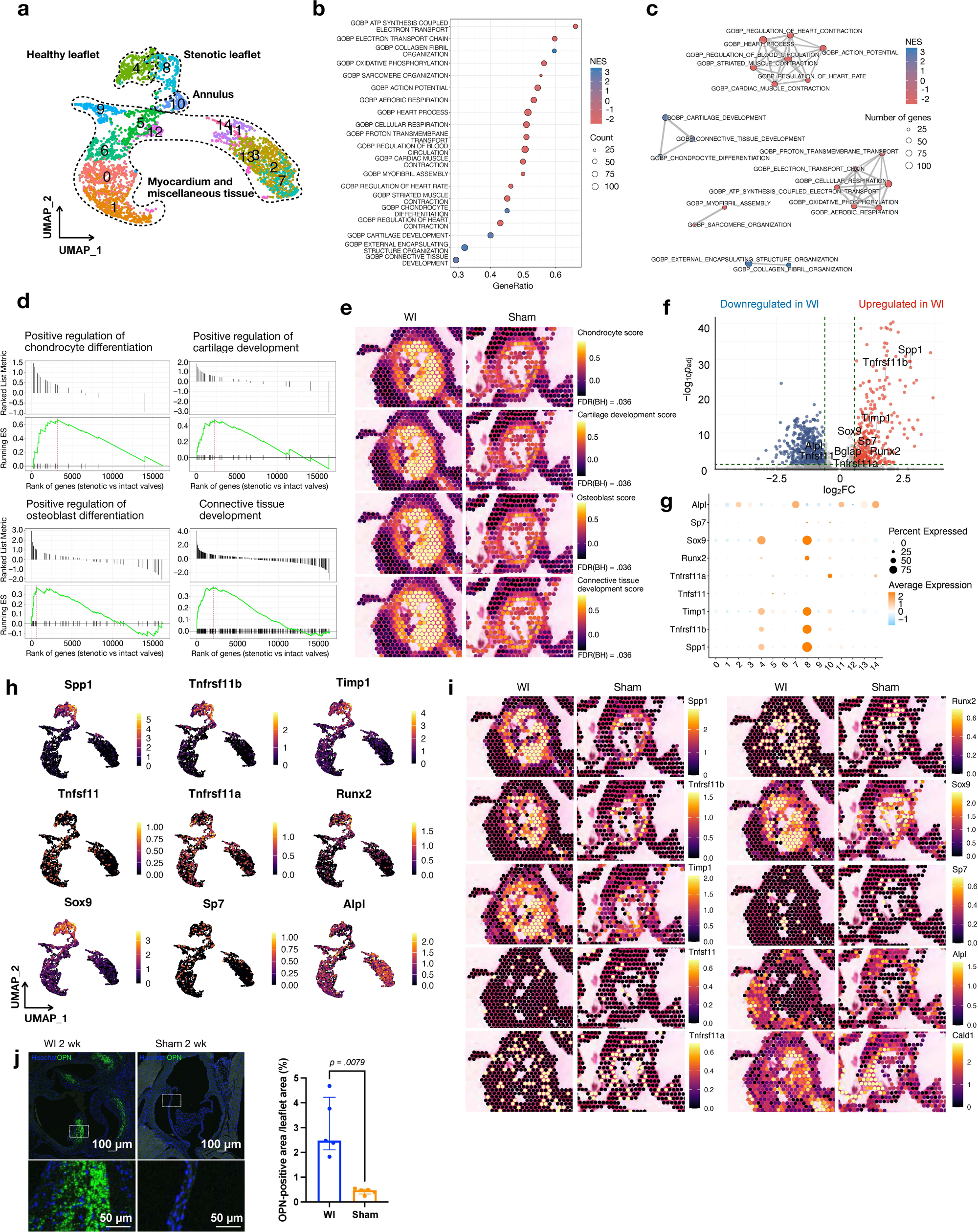
Early osteochondrogenic gene programs emerge in valve leaflets following endothelial injury. **a**, UMAP of merged Visium data from wire injury (WI) and sham valves (*n* = 3 per group). **b–c,** GSEA of stenotic versus healthy leaflets (GO Biological Process). **b,** Bubble plot: GeneRatio (x), gene count (size), and normalized enrichment score (NES; blue-enriched, red-depleted). **c,** Enrichment map: node size = count, color = NES, edge = gene set overlap. **d,** GSEA enrichment plots for representative osteochondrogenic GO terms in stenotic versus healthy leaflet regions: positive regulation of chondrocyte differentiation, cartilage development, osteoblast differentiation, and connective tissue development. Green curves show the running enrichment score, tick marks indicate gene hits, and positive NES indicates enrichment in the stenotic leaflet. **e**, Spatial module scores for osteochondrogenic programs (positive regulation of chondrocyte differentiation, positive regulation of cartilage development, positive regulation of osteoblast differentiation, and connective tissue development). Right: Module score and Benjamini–Hochberg (BH) false discovery rate (FDR) values for each gene set. **f**, Differential expression between stenotic and healthy leaflets. x-axis, log2 fold-change (stenotic/healthy); y-axis, −log10 Bonferroni-adjusted *p*. Red, genes upregulated in stenotic regions; blue, genes downregulated in stenotic regions; grey, not significant. Dashed lines indicate thresholds (|log2FC| > 0.585, corresponding to a fold change > 1.5, and Bonferroni-adjusted *p* < 0.05). Selected osteochondrogenic genes were annotated. **g,** Osteochondrogenic markers across Visium clusters (0–14. Dot size indicates the percentage of spots expressed, and color denotes the scaled average expression. Key clusters: 4, healthy leaflet tissues; 8, stenotic leaflet tissues; and 10, annulus. **h–i,** Feature plots of osteochondrogenic markers in the aortic valve and adjacent short-axis tissues for the WI and sham groups. **h,** UMAP with expression overlays from Visium spots; the color indicates the expression level (scale on the right). The cluster identities match Fig. 2a a. **i,** Spatial expression maps; color scale on the right. **j**, Left, IF for Osteopontin (OPN; *Spp1*) in the aortic valves from WI and sham groups (scale bars: 100 µm, top; 50 µm, bottom); right, quantification of the OPN^+^ DAB area fraction (%) by IHC. n = 5 mice per group. Data are presented as median (IQR), assessed using two-sided Mann–Whitney *U* tests.

To identify disease-associated programs in early-stage AS, we performed gene set enrichment analysis (GSEA; Gene Ontology (GO) Biological Process (BP)) to compare stenotic and intact leaflets. Enrichment was quantified using a normalized enrichment score (NES). The top-ranked terms by gene ratio indicated the depletion of mitochondrial respiration processes (ATP synthesis-coupled electron transport, electron transport chain, and oxidative phosphorylation; negative NES) and the enrichment of connective tissue and chondro/osteogenic development, including connective tissue development, collagen fibril organization, cartilage development, and chondrocyte differentiation (positive NES) (Fig. 2b). Significant gene sets (false discovery rate (FDR) < 0.05) were organized into similarity-based modules in an enrichment map, highlighting mitochondrial/respiratory, muscle contraction, and matrix/chondro-osteogenic programs (Fig. 2c).

Running enrichment plots for the positive regulation of chondrocyte differentiation, positive regulation of osteoblast differentiation, and connective tissue development showed positive NES values with early accumulation of leading-edge genes, supporting the activation of osteochondrogenic programs at this precalcific stage (Fig. 2d; Supplementary Data 1).

Spatial feature plots displayed four chondro-osteogenic module scores comprising GOBP Chondrocyte differentiation (chondrocyte score), GOBP Cartilage development (cartilage development score), GOBP Osteoblast differentiation (osteoblast score), and GOBP Connective tissue development (Connective tissue development score), which were elevated in thickened leaflets relative to morphologically intact leaflets (FDR = 0.036 for all; Fig. 2e; Supplementary Data 1).

Spatial enrichment mapping demonstrated that these pathway signatures were predominantly enriched within the interstitial regions of stenotic valve leaflets (Fig. 2e). Bonferroni-adjusted differential gene expression (DGE) analysis of calcification- and matrix-remodeling-associated genes (17–22) showed upregulation of *Spp1*, *Tnfrsf11b*, *Timp1*, *Runx2*, and *Sox9* in thickened leaflets versus intact valves, whereas *Tnfrsf11a*, *Tnfsf11*, *Sp7*, and *Alpl* showed no significant changes (Fig. 2f). Dot plot demonstrated increased fractions of expressed spots together with higher mean expression levels of osteogenic and chondrogenic genes in Cluster 8 than in Cluster 4 (Fig. 2g). Consistently, violin plots showed right-shifted expression distributions for the upregulated genes, further supporting activation of early osteochondrogenic programs in thickened valve leaflets (Supplementary Fig. 4a); a heatmap summarized the marker set (Supplementary Fig. 4b). Feature plots of UMAP embedding suggested the enrichment of osteochondrogenic genes in cluster 8 (Fig. 2h), and spatial feature plots of these genes were placed as high-expression spots in the corresponding thickened leaflet regions in tissue coordinates (Fig. 2i). Orthogonal validation by IF and chromogenic IHC confirmed increased osteopontin (OPN/SPP1); the OPN^+^ area fraction was significantly larger in WI mice than in sham mice (Fig. 2j; *p* = 0.0079), consistent with the transcriptomic results. These findings indicate that selective osteochondrogenic transcriptional programs are activated in thickened valve leaflets during the precalcific stage.

### Leaflet thickening is predominantly driven by *PDGFRβ*^+^ valvular interstitial cells

Next, to identify the cellular source driving leaflet thickening, a major contributor to AS, we focused on the EndMT signaling pathway because accumulating evidence indicates that inflammatory cytokines and disturbed flow can promote EndMT in the valvular endothelium (23–26). Because no dedicated EndMT gene set was available in the GO Biological Process or MSigDB Hallmark collections, we used the hallmark epithelial–mesenchymal transition (EMT) gene set as a pragmatic surrogate and supplemented it with literature-curated mesenchymal gain and endothelial-identity marker sets (27–32) to calculate module scores (Fig. 3a; Supplementary Data 1, 2). In the GSEA plot, the running enrichment score peaked early with a left-skewed barcode, indicating positive enrichment of EMT genes in the thickened valvular regions (Fig. 3b). To assess early EndMT-like transition, we examined a targeted marker panel encompassing endothelial identity (*Pecam1*, *Cdh5*, *Klf2*, *Cldn5*, *Klf4*, *Tek*, *Kdr*, and *Emcn*) and EMT/mesenchymal activation (*Snai1*, *Snai2*, *Postn*, *Taglin*, *Acta2*, and *Meox1*). In injured valve cluster 8 versus morphologically intact cluster 4, *Pecam1*, *Klf2*, *Kdr*, and *Tek* were significantly decreased, whereas *Snai1*, *Snai2*, *Tagln*, and *Acta2* were increased (Bonferroni-adjusted *p* < 0.05); *Meox1*, *Cdh5*, *Cldn5*, and *Emcn* showed no significant change (Fig. 3c). Violin and dot plots across Visium clusters 1–14 visualized the same nine-gene panel used in the DGE analysis, with cluster-wise patterns concordant with those results. The injured valve cluster 8 showed reduced *Pecam1*, with relative stability of *Cdh5*, *Meox1*, *Cldn5*, and *Emcn* alongside increased *Snai1*, *Snai2*, *Tagln*, and *Acta2*, compared with the morphologically intact cluster 4 (Fig. 3d, e). A heatmap of the same 12-gene panel across the Visium spots showed that unsupervised hierarchical clustering separated endothelial-identity genes from mesenchymal and EMT markers. The injured valve cluster 8 displayed a mesenchymal-biased pattern with attenuated *Pecam1*, whereas the morphologically intact cluster 4 showed a reciprocal profile (Fig. 3f). UMAP embeddings with Visium spot-level overlays and spatial feature maps of the same eight-gene panel corroborated the preceding analyses, which showed mesenchymal gain with attenuated endothelial identity that localized to stenotic leaflets in cluster 8, whereas the morphologically intact cluster retained endothelial signals (Fig. 3g, h).

**Fig. 3.**
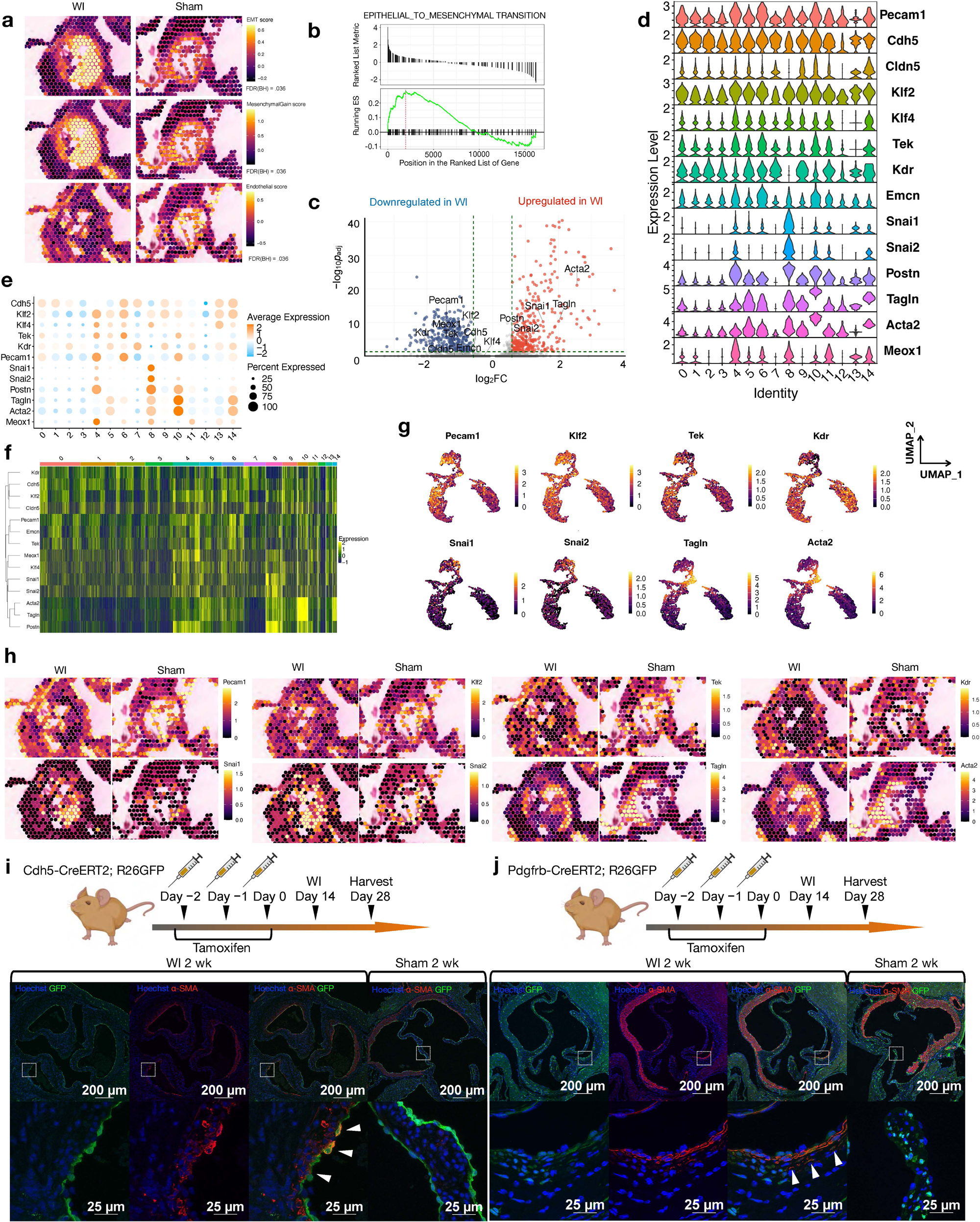
Endothelial-to-mesenchymal transition programs localize to regions bearing osteochondrogenic gene signatures, supported by lineage tracing. a, Spatial feature plots of module scores for GOBP term epithelial-to-mesenchymal transition and EndMT-associated signatures, including literature-curated mesenchymal gain and endothelial cell scores. Right, module score and FDR values for each gene set. **b,** GSEA enrichment plot for the GOBP term epithelial-to-mesenchymal transition, comparing stenotic and healthy leaflet regions; the green curve shows the running enrichment score, and ticks denote gene hits; positive NES indicates enrichment in stenotic leaflets. **c,** Volcano plot of differentially expressed genes between stenotic and healthy leaflets. x-axis: log2 FC (stenotic/healthy); y-axis: −log10 Bonferroni-adjusted *p*. Red, genes upregulated in stenotic regions; blue, genes downregulated in stenotic regions; grey, not significant. Dashed lines indicate thresholds (|log2FC| > 0.585, corresponding to a fold change > 1.5, and Bonferroni-adjusted *p* < 0.05). Selected mesenchymal and endothelial markers were annotated. **d**, Violin plots showing normalized expression per cluster for the same gene panel; cluster identities are shown in Figure 2a. **e,** Dot plot of endothelial and mesenchymal markers across visible clusters (0–14. Dot size indicates the percentage of spots expressed, and color denotes the scaled average expression. Key clusters: cluster 4, healthy leaflet tissue; cluster 8, stenotic leaflet tissue; and cluster 10, annulus. UMAP embeddings with visible-spot-level expression overlays for the same panel; the color indicates the expression levels (scale shown). **f,** Heatmap of the same nine-gene panel across Visium spots (clusters 0–14); color denotes gene-wise scaled expression (z-score). A row dendrogram shows the unsupervised hierarchical clustering of genes, and the columns are grouped by cluster identity, as shown in Fig. 2a. **g–h,** Feature plots of endothelial- and mesenchymal-related markers. **g**, UMAP with Visium spot-level expression overlays; the color indicates normalized expression (scale shown). The cluster identities are shown in Figure 2a. **h**, Spatial expression maps across Visium sections encompassing the aortic valve and neighboring myocardium. Color indicates normalized expression (scale shown). **i–j**, Lineage tracing in tamoxifen-inducible Cdh5-CreERT2;R26GFP and Pdgfrb-CreERT2;R26GFP mice in which endothelial (**i**) or valvular interstitial cells (**j**) and their descendants express GFP. White arrowheads show α-SMA and GFP double-positive cells. WI, wire injury; wk, week.

To confirm whether VECs and their progeny contribute to early-stage valve thickening, we performed lineage tracing of VECs using tamoxifen-inducible Cdh5-CreER^T2^-driven reporter mice (Cdh5-CreER^T2^; ROSA EGFP mice), in which VECs and their descendants were permanently labeled with GFP. Following the WI procedure, injured VECs co-expressing GFP and α-SMA localized along the endothelial lining of the valve leaflets, whereas α-SMA^+^ cells adjacent to the endothelium were negative for GFP, which indicated that VECs and their descendants were not observed in the valvular interstitial space (Fig. 3i). Notably, disrupted α-SMA^+^ endothelial cell–cell interactions were also observed in WI reporter mice (Fig. 3i, white arrowheads). Next, we focused on mesenchymal cellular markers, PDGF receptors, and performed WI on tamoxifen-inducible Pdgfrb-CreER^T2^ reporter mice (Pdgfrb-CreER^T2^; ROSA EGFP mice), in which VICs and their descendants were permanently labeled with GFP. In the valvular interstitial space, α-SMA^+^GFP^+^ VICs were observed (Fig. 3j), whereas VECs did not show GFP labeling (Fig. 3i). Additionally, GFP^+^ VICs co-expressed heat shock protein 47 (HSP47), a collagen-specific chaperone known as a marker of collagen biosynthesis and maturation, indicating that profibrotic programs were induced in PDGFRβ^+^ VICs following endothelial injury (Supplementary Fig. 5). These findings indicate that PDGFRβ^+^ VICs undergo profibrotic activation following endothelial injury and constitute a major cellular source contributing to leaflet thickening.

### Endothelial injury activates the TGF-β–CTHRC1 axis in the valvular interstitium

To investigate signaling pathways involved in tissue remodeling following VEC injury, we examined whether TGF-β signaling, a key inducer of collagen production, is activated in valve leaflets.

Representative tissue-remodeling genes associated with profibrotic and myofibroblast-like programs, including *Tgfb1*, *Tgfbi*, *Col1a1*, *Col3a1*, *Acta2* (α-SMA), *Serpinh1* (HSP47), *Lox*, *Fn1*, *Cthrc1*, and *Postn* (33–40), were upregulated in stenotic valvular tissues (Fig. 4a). Dot plot of TGF-β signaling– related genes revealed a greater proportion of expressing spots together with higher mean expression levels in cluster 8 than in cluster 4 (Fig. 4b). Consistently, violin plots demonstrated right-shifted expression distributions of these profibrotic genes in stenotic valvular tissues (Supplementary Fig. 6a). The UMAP feature plots localized the fibrosis program enrichment to cluster 8, with *Acta2* marking a subpopulation transcriptionally adjacent to the annulus (cluster 10), indicating an interface between leaflet myofibroblast activation and annular compartments at this early time point (Supplementary Fig. 6b). The heatmap and hierarchical clustering resolved two major co-expression branches: an upper contractile/remodeling branch containing *Acta2*, *Lox*, and *Cthrc1* and a broader matrix/fibrotic branch containing *Pdgfrb*, *Serpinh1*, *Col3a1*, *Col1a1*, *Fn1*, *Tgfb1*, and *Tgfbi*. Notably, *Cthrc1* clustered most closely with *Lox* and was markedly upregulated in the diseased leaflet tissue cluster 8 relative to the intact leaflet tissue cluster 4 (Supplementary Fig. 6c).

**Fig. 4.**
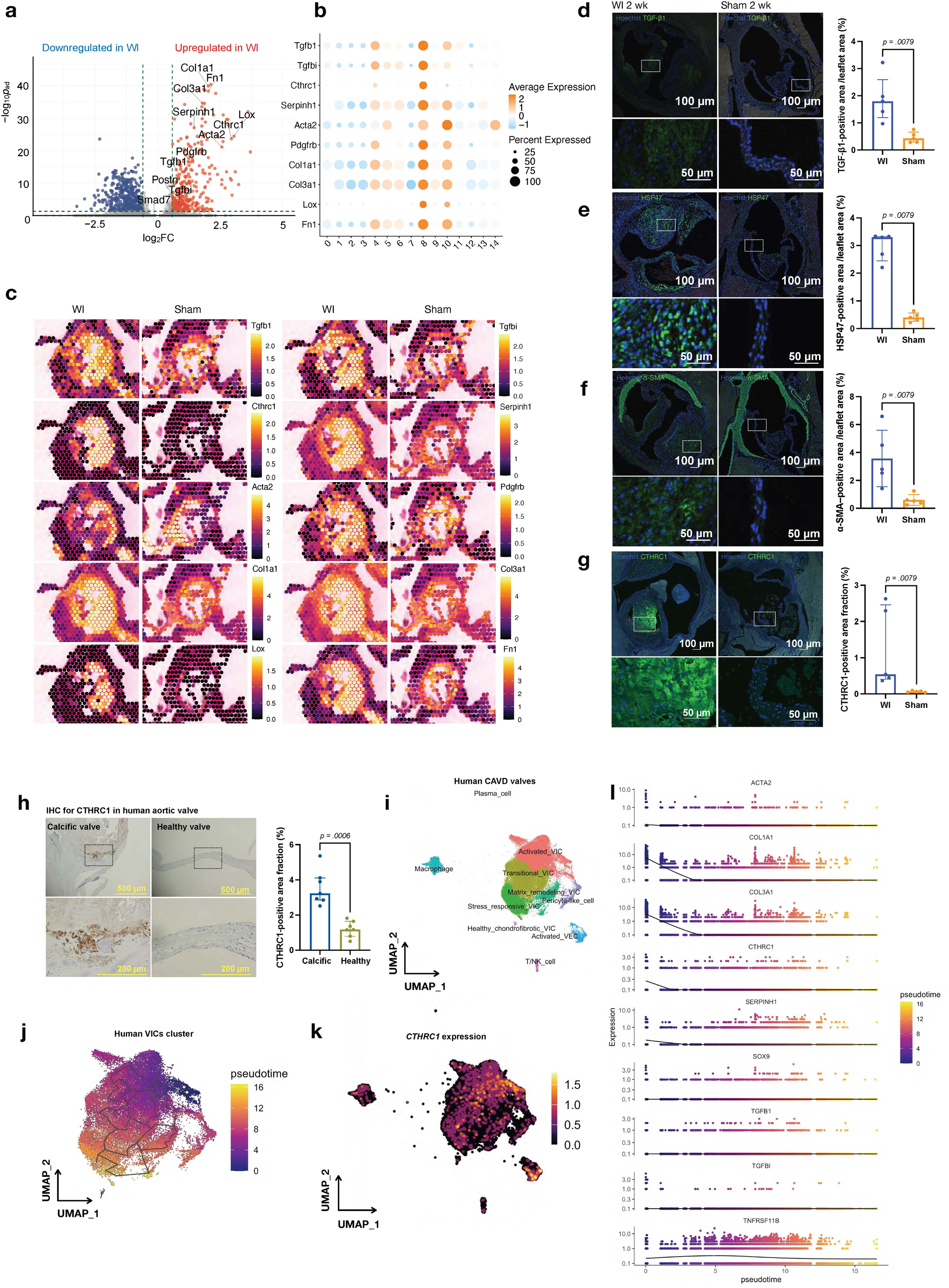
Endothelial injury enhances TGF-β1 and downstream profibrotic signaling in the valvular interstitium. **a**, Volcano plot for stenotic versus healthy leaflets: x-axis, log2FC (stenotic/healthy); y-axis, −log10 Bonferroni-adjusted *p*. Selected fibrosis/remodeling associated genes are labeled. **b–c,** Expression summaries for fibrosis/remodeling-associated genes across Visium clusters (0–14. **b**, Dot plot (dot size = percentage of spots; color = scaled average expression). **c**, Spatial expression maps; color represents expression levels (right scale). **d–g**, Left, IF of aortic valves from wire injury (WI) and sham mice; right, IHC quantification of the DAB^+^ area fraction (DAB^+^ area/leaflet area, %). Proteins: **d**, TGF-β1; **e**, HSP47; **f**, α-SMA; **g**, CTHRC1. Scale bars: 100 µm (top), 50 µm (bottom). n = 5 mice per group. Data are presented as median (IQR) and assessed using two-sided Mann–Whitney *U* tests. **h,** Representative IHC images (left) and quantification of CTHRC1^+^ DAB area fractions (right) in human calcific aortic valve disease (CAVD) valves and healthy control valves obtained from cadaveric donors. Scale bars, 500 μm (top) and 200 μm (bottom). n = 7 donors per group. Data are presented as median (IQR) and assessed using two-sided Mann–Whitney *U* tests. **i–l,** Re-analysis of publicly available scRNA-seq data from human CAVD and healthy valves. **i,** UMAP visualization of the annotated cell populations. **j**, Pseudotime analysis of valvular interstitial cells (VICs). **k,** Feature plot showing *CTHRC1* expression. **l**, Pseudotime analysis of fibrotic-remodeling programs in VICs. wk, week.

Spatial feature plots illustrated focal enrichment of *Acta2* in the annulus- and endothelium-adjacent thickened leaflets, whereas other representative fibrosis markers were diffusely upregulated across the thickened leaflet compartment (Fig. 4c). IF analysis demonstrated markedly increased signals for TGF-β1, HSP47, α-SMA, and CTHRC1 within thickened regions of injured valve leaflets during the early stage after WI, consistent with focal activation of profibrotic remodeling pathways.

Complementary chromogenic immunohistochemistry further confirmed these findings, as the quantified DAB^+^ area fractions determined by color deconvolution were significantly greater in the WI group than in the sham group (Fig. 4d–g). Because DGE analysis identified *Cthrc1* among the most significantly enriched genes in thickened valve leaflets, and prior studies have implicated CTHRC1 as a potential therapeutic regulator in AS (41,42), we next examined its functional role in valvular remodeling and calcification. Immunohistochemical analysis of human AS/CAVD specimens demonstrated prominent CTHRC1 deposition. Quantification of the CTHRC1^+^ area fraction confirmed that this remodeling-associated protein was enriched in human diseased valves (Fig. 4h). In addition, we reanalyzed a previously published scRNA-seq dataset of human calcific aortic valves available under the NCBI BioProject accession PRJNA562645 (25). Following batch-effect correction, integration, and clustering, the VICs were further classified into activated, transitional, pericyte-like, matrix-remodeling, and stress-response clusters (Fig. 4i; Supplementary Fig. 7a).

Pseudotime analysis of the human transcriptomic dataset revealed that the activated VIC subcluster, corresponding to the early stage of CAVD, exhibited high expression of *CTHRC1* (Fig. 4j–l; Supplementary Fig. 7b). These findings are consistent with our independent observations in human valve specimens from IF/IHC analyses, as well as with murine transcriptomic and protein expression analyses.

### CTHRC1 protects against valvular calcification by limiting foamy macrophage infiltration

We next performed the WI procedure in *Cthrc1^−/−^* mice to compare pathological changes between the wild type and *Cthrc1^−/−^* groups after endothelial injury to the murine aortic valves. A two-way analysis of variance (ANOVA) of serial echocardiographic measurements demonstrated that although the AV peak velocity was transiently elevated in knockout mice during the acute phase within 3 days after WI, no significant differences were observed between the groups at 1 and 2 weeks after injury (Fig. 5a, Supplementary Fig. 8), indicating no clear effect of *Cthrc1* status on the hemodynamic severity of AS during the early stage. As hemodynamics of AS in the *Cthrc1^−/−^* group were comparable to those of wild type WI group in early-stage AS, we therefore focused on chronic-phase pathological features for subsequent analyses. At 16 weeks after WI, von Kossa staining revealed an increased von Kossa^+^ area fraction in *Cthrc1^−/−^* mice compared with wild-type controls (Fig. 5b), suggesting that loss of CTHRC1 is associated with enhanced valvular calcification in the chronic phase.

**Fig. 5.**
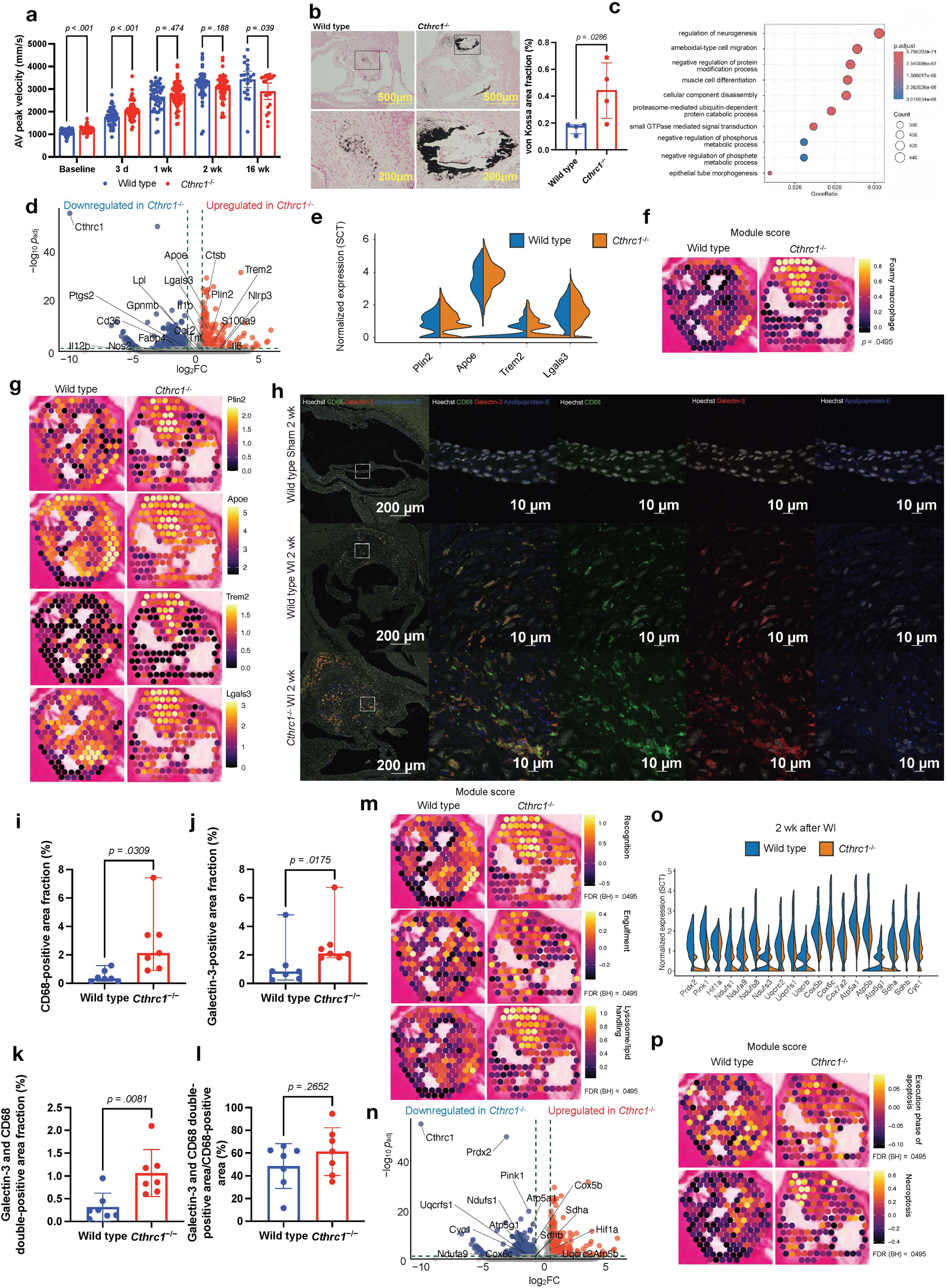
Galectin-3^+^ foamy macrophage-associated signatures are enriched in stenotic aortic valves, and apoptosis-related signatures are prominent under CTHRC1-deficient conditions. **a**, Serial AV peak velocity in wild-type and *Cthrc1^−/−^* mice after wire injury (WI); sample sizes per time point (n, wild type/Cthrc1^−/−^): baseline, 64/91; 3 days (d), 55/62; 1 week (wk), 49/71; 2 wk, 49/57; 16 wk, 21/20. **b**, Left, representative von Kossa-stained images of the aortic valves from wild-type and *Cthrc1^−/−^* mice 16 weeks after WI (scale bars, 500 µm, top; 200 µm, bottom); right, quantification of the von Kossa^+^ area fraction (%). n = 4 mice per group. Data are presented as median (IQR) and assessed using two-sided Mann–Whitney *U* tests. **c**, The top 10 GO terms ranked by GeneRatio from enrichment analysis comparing *Cthrc1^−/−^* and wild-type stenotic valves. X-axis, gene ratio; dot size, gene count; color, adjusted-*p.* d, Differential expression of *Cthrc1^−/−^* versus wild-type stenotic leaflet: x-axis, log2FC; y-axis, −log10 Bonferroni-adjusted *p*. Selected macrophage-associated genes are labeled. e, *Plin2*, *Apoe, Trem2,* and *Lgals3* expression in the two groups. **f,** Spatial feature plots of module scores for literature-curated foamy macrophage signatures. Right, module score and FDR for each gene set. **g,** Spatial expression maps. Right: expression levels. **h**, Representative multichannel IF images of aortic valves from wild-type sham mice (top), wild-type mice 2 wk after WI (middle), and *Cthrc1^−/−^* mice 2 wk after WI (bottom). Left, low-magnification images (scale bars, 200 µm); remaining panels, high-magnification images (scale bars, 10 µm). **i–l**, Quantification of CD68^+^ (**i**), Galectin-3^+^ (**j**), CD68^+^ Galectin-3^+^ area fraction (**k**), and CD68^+^ Galectin-3^+^ /CD68^+^ leaflet area (**l**). n = 7 mice per group. Data are mean ± s.d. or median (IQR); Welch’s t-test or two-sided Mann–Whitney *U* test. **m,** Module scores for literature-curated gene sets related to opsonin/bridging, engulfment, and lysosomal lipid handling. Right, module scores and FDR values for each gene set. **n**, Volcano plot. Selected mitochondrial and respiratory chain-associated genes were labeled. **o**, Violin plots comparing the expression levels of representative mitochondrial/respiratory-associated genes between the two groups. **p,** Module scores for literature-curated gene sets related to the GO term execution phase of apoptosis and the literature-curated necroptosis score. Right, module scores and FDR values for each gene set.

To characterize the early transcriptional changes associated with enhanced valvular calcification observed in *Cthrc1*-deficient mice, we performed spatial transcriptomic analysis 2 weeks after WI. We restricted the analysis to the aortic valve leaflet region, which was manually annotated in the Loupe Browser (v7.0.1). After a quality check of the specimens (Supplementary Fig. 9a, b), GO enrichment analysis identified amoeboidal-type cell migration (GO:0001667) as a significantly enriched term (Fig. 5c). In line with this finding, we examined macrophage infiltration into the valvular tissue. DGE analysis revealed a marked upregulation of *Trem2*, *Plin2*, *Lgals3*, and *Apoe*, which are preferentially expressed by foamy macrophages (43–45), whereas genes associated with inflammatory and classically activated myeloid states, including *Il1b*, *Nos2*, *Il12b*, *Tnf*, and *Ccr2* (46,47), were not significantly upregulated (Fig. 5d). In the *Cthrc1^−/−^* group, *Plin2*, *Apoe, Trem2,* and *Lgals3* displayed an increased proportion of high-expression spots relative to the wild-type control, reflected by a heavier upper tail in the expression distributions (Fig. 5e). Consistent with these findings, spatial feature mapping revealed a higher module score of foamy macrophages in stenotic aortic valves from *Cthrc1^−/−^* mice (*p* = 0.0495; Fig. 5f, Supplementary Data 3) and increased expression of *Plin2*, *Apoe, Trem2,* and *Lgals3* within thickened valves compared with those in wild-type controls (Fig. 5g). Protein-level validation by IF staining demonstrated increased area fractions positive for Galectin-3, positive for CD68, and double-positive for Galectin-3 and CD68 within the thickened valves in *Cthrc1^−/−^* group compared with the wild-type group (Fig. 5h–k). By contrast, the Galectin-3/CD68 double-positive area, normalized to the CD68^+^ area, was not significantly increased in the *Cthrc1^−/−^* group relative to the wild-type group, suggesting similar profiles of migrating foamy macrophages between the two groups (Fig. 5l).

To further characterize the foamy macrophage-associated transcriptional landscape in lesions from *Cthrc1^−/−^* mice, in which Galectin-3-rich areas were observed, we performed module score analysis using custom gene sets related to opsonin/bridging (displayed as “recognition” in the figures), engulfment, and lysosome/lipid handling (Supplementary Data 3). All three module scores were significantly increased (FDR (Benjamini–Hochberg [BH]) = 0.0495 for each module; Fig. 5m), consistent with enhanced lipid handling in thickened valvular lesions under *Cthrc1*-depleted conditions.

In addition to these enrichments, we observed increased *Hif1a* expression and decreased *Prdx2* expression together with reduced expression of genes encoding components of the canonical mitochondrial oxidative phosphorylation machinery, including *Uqcrfs1*, *Cyc1*, *Cox5b*, *Atp5a1*, and *Atp5g1*, in the *Cthrc1^−/−^* mice (Fig. 5n, o). Moreover, the module scores for GO:0097194 (execution phase of apoptosis) and the necroptosis signature were modestly but significantly higher in *Cthrc1^−/−^* mice than in wild-type controls (Fig. 5p; Supplementary Data 3), indicating a shift toward activation of apoptotic execution and necroptosis-related programs in *Cthrc1^−/−^* valves. Overall, our data show coordinated alterations in mitochondrial respiratory chain genes, increased apoptosis- and necroptosis-related programs, and enrichment of foamy cell-associated macrophage signatures in stenotic valvular lesions in the *Cthrc1^−/−^*group.

## Discussion

In this study, we established an optimized murine model of valvular endothelial injury that rapidly recapitulates the key hemodynamic and histopathological features of AS/CAVD. Spatial transcriptomic profiling during the acute phase following injury revealed the early activation of osteochondrogenic and profibrotic transcriptional programs within the thickened valve leaflets before the development of overt calcification. Lineage-tracing analyses demonstrated that endothelial cells undergoing mesenchymal transition–associated changes remained largely confined to the leaflet surface, whereas VICs with high PDGFRβ promoter activity represented the predominant cellular source contributing to leaflet thickening. We further identified activation of the TGF-β1–CTHRC1 axis in injured valvular tissue. Functional analyses demonstrated that *Cthrc1* deficiency exacerbated valvular calcification and promoted the accumulation of foamy macrophages without broadly altering fibrotic transcriptional programs, suggesting that CTHRC1 may protect against maladaptive inflammatory remodeling during AS progression.

Prevailing models of early-stage AS/CAVD typically emphasize valvular endothelial dysfunction, the Lp(a)-OxPL-autotaxin-lysophosphatidic acid axis, immune-inflammatory circuits, osteogenic reprogramming driven by the BMP2-RUNX2/Wnt pathways, mechanotransduction programs such as Piezo1-YAP, and extracellular vesicle-mediated nucleation as key processes in CAVD (13,48–50), yet directly tracking disease onset *in situ* remains technically challenging. We used a murine model that permits the precise control of harvest timing, enabling stage-resolved sampling across disease progression. Because end-stage AS/CAVD can be investigated using human specimens, we focused on the earliest stages, which we operationally defined as a lack of microscopic evidence of calcification. Activated VECs and VICs expressing myofibroblastic markers were identified in early-stage murine AS/CAVD specimens. Several scRNA-seq studies have reported EndMT-related findings (6,7,25) and the potential relocation of VECs toward the valvular interstitium in this context (51), and our spatial transcriptomic data were also compatible with the activation of VECs and VICs. However, because the transcriptional state alone does not resolve the spatial position or cell movement, these observations cannot distinguish physical migration from local phenotypic remodeling. In our lineage-tracing analysis of early lesions, we found no evidence of overt displacement of activated VECs into the leaflet interstitium. Instead, VECs upregulated α-SMA, adopted a contracted morphology, and showed disrupted endothelial barrier integrity. These findings support a model in which endothelial plasticity at the leaflet surface initiates interstitial remodeling without requiring endothelial emigration. Consistently, EndMT-associated regions showed an enrichment of transcriptional programs related to mononuclear cell migration, together with increased immune-cell-associated signatures, including monocytes, plasma cells, and foamy/lipid-associated macrophages (Fig. 1j, Supplementary Fig. 9a–g). Consistent with these transcriptomic inferences, immunostaining identified CD68^+^ cells and CD68^+^ cellular remnants within the valvular interstitium at the earliest post-onset time point examined (Fig. 1j). Together, these data recapitulate the inflammatory component emphasized in prior reviews and scRNA-seq data of human CAVD specimens (25) (Supplementary Fig. 10a–d) and support the early recruitment of mononuclear phagocytes to the affected tissue (7,13,52). Interestingly, our transcriptomic data indicated that macrophage response to WI was qualitatively reshaped by *Cthrc1* deficiency. In wild-type mice, WI did not induce a detectable upregulation of *Plin2*, *Trem2*, or *Apoe*, which are widely used markers of foamy macrophages (43–45), relative to sham controls (Supplementary Fig. 11a–g). By contrast, when both groups underwent the same WI under normal chow conditions, these transcripts and *Lgals3* were robustly upregulated in *Cthrc1^−/−^* mice compared with wild-type counterparts. Collectively, these findings indicate that the loss of *Cthrc1* amplifies a foamy macrophage-like transcriptional program in response to endothelial ablation and subsequent serum component entry into the valvular interstitium (Fig. 6).

**Fig. 6.**
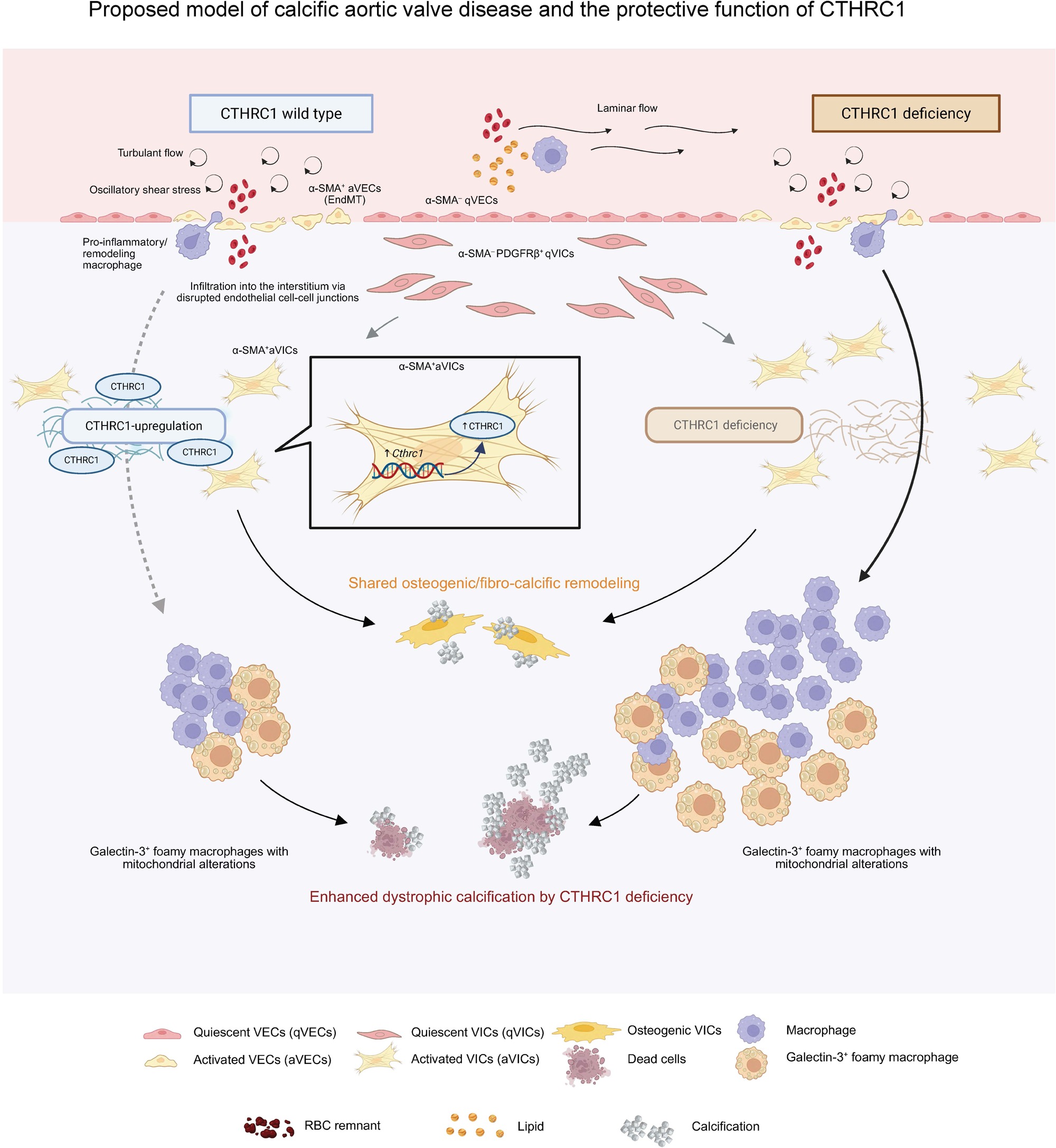
Proposed model of calcific aortic valve disease progression and the protective role of CTHRC1. Proposed model of calcific aortic valve disease (left side). RBCs and immune cells, including CD68^+^ macrophages, enter the valvular interstitium from the circulation following endothelial injury and activation. Although CTHRC1-mediated interstitial remodeling occurs, Galectin-3^+^ foamy macrophages persist in the valvular interstitium. In contrast, under CTHRC1-deficient conditions, CD68^+^ Galectin-3^+^ cells accumulate in the aortic valve interstitium after endothelial injury and activation, accompanied by the downregulation of mitochondrial respiratory chain genes and increased apoptosis- and necroptosis-related programs. Although the osteogenic calcification signatures remained comparable to those in CTHRC1-preserved conditions, valvular interstitial calcification was significantly exacerbated. These findings support a model in which CTHRC1 limits dystrophic calcification-associated remodeling rather than osteogenic calcification by regulating foamy macrophage accumulation in stenotic aortic valves.

CTHRC1 functions as a context-dependent regulator of tissue remodeling downstream of TGF-β1, where it limits pathway duration by facilitating proteasomal turnover of pSmad2/3. In addition, CTHRC1, a secreted protein implicated in tissue/matrix remodeling (53,54), is induced after endothelial injury in the aortic valve (55) and is enriched in calcific disease-associated valvular interstitial cell populations (41,42). In our dataset, the expression levels of *Tgfb1*, its downstream target genes, and collagen-associated genes in *Cthrc1^−/−^* AS valves were not significantly altered compared with those in wild-type AS valves (Supplementary Fig. 12a), suggesting that exaggerated canonical fibrotic signaling is unlikely to fully explain the enhanced calcification phenotype observed under *Cthrc1* deficiency. In contrast, foamy macrophage infiltration and associated lipid handling programs are enhanced in stenotic valves lacking *Cthrc1*. These findings raise the possibility that CTHRC1 may inhibit inflammatory macrophage accumulation independent of major transcriptional changes in fibrotic remodeling pathways. Given that dystrophic calcification is closely linked to chronic inflammatory cell accumulation and defective tissue resolution, the enhanced persistence of foamy macrophages may contribute to the accelerated calcific remodeling observed in *Cthrc1*-deficient valves. Although the precise mechanism remains unclear, CTHRC1 may influence inflammatory cell retention, extracellular matrix organization, and lipid-associated microenvironmental remodeling within injured valvular tissue.

In our study, *Cthrc1* deficiency aggravated valvular calcification in an AS/CAVD model. However, relative to wild-type lesions, *Cthrc1*-deficient lesions did not exhibit significantly higher module scores for GO terms related to osteoblast differentiation, chondrocyte differentiation, or chondrocyte development (Supplementary Fig. 12b and c). In contrast, mitochondrial dysfunction and cell death-related programs, including the GO-based execution phase of apoptosis and a literature-curated necroptosis signature, were significantly enriched in *Cthrc1*-deficient lesions (Fig. 5p). Interestingly, mitochondrial dysfunction- and respiratory chain-related signatures were downregulated in the wild-type WI group compared with those in the sham group (Supplementary Fig. 13a–g), and this downregulation was further exacerbated in the *Cthrc1*-deficient group. These findings suggest that the enhanced calcification observed in *Cthrc1*-deficient valves is not accompanied by a detectable increase in canonical osteogenic or chondrogenic transcriptional programs at the analyzed stage but is instead associated with transcriptional signatures of mitochondrial dysfunction and cell death-related injury responses. Taken together, AS/CAVD exhibits a shared osteogenic/fibrocalcific remodeling program, whereas *Cthrc1* deficiency appears to selectively exacerbate degenerative remodeling characterized by foamy macrophage accumulation, mitochondrial dysfunction, apoptosis, necroptosis-related signatures, and dystrophic calcification, thereby increasing overall valve calcification. The concomitant increase in apoptosis- and necroptosis-related scores further supports the transition toward a lesion state permissive for mineral deposition, as regulated cell death pathways are increasingly being recognized as contributors to calcification in cardiovascular tissues. Although our data did not establish a direct causality between these transcriptional alterations and late mineral accumulation, the increased von Kossa^+^ area in the chronic phase is consistent with a model in which macrophage accumulation, metabolic stress, and cell death programs cooperate to accelerate calcific remodeling (56,57) in *Cthrc1*-deficient valves (Fig. 6). Further pathway analysis using Ingenuity Pathway Analysis (IPA) of injured wild-type and *Cthrc1*-deficient valves revealed enhanced macrophage activation, whereas pathways associated with tissue remodeling and fibrosis, including collagen production, were not significantly altered. These findings suggest that *Cthrc1* is unlikely to regulate extracellular matrix remodeling to limit myeloid cell infiltration but may instead directly modulate the activation and function of infiltrating myeloid cells (Supplementary Fig. 14). Canonical pathway analysis further identified the enrichment of neutrophil degranulation, antigen presentation, interferon signaling, and ferroptosis signaling pathways in *the Cthrc1*-deficient valves (Supplementary Fig. 15). Notably, TFEB-mediated phagocytosis was more strongly activated in the *Cthrc1*-deficient valves than in the wild-type valves (Supplementary Fig. 16). Given that interferon signaling is a key driver of M1-like inflammatory macrophage polarization and that CTHRC1 promotes the phenotypic transition of macrophages from an M1-like inflammatory state to an M2-like anti-inflammatory state in colorectal cancer (58), our findings raise the possibility that CTHRC1 suppresses inflammatory phagocyte activation, thereby limiting calcification associated with cell death.

This study has several limitations. First, our conclusions were based on an animal model, and the tempo and magnitude of disease evolution may differ from the protracted course observed in humans, potentially influencing the timing and dynamic range of early pathological phenotypes. Second, formal genetic rescue of the late-stage phenotype is technically challenging because disease progression involves irreversible tissue remodeling and calcification, thereby limiting the feasibility of within-animal reversal once the pathology is established. Third, the genetically modified mouse lines used in this study were maintained on different genetic backgrounds. In particular, *Cthrc1^−/−^* mice were maintained on a mixed genetic background, whereas the wild-type control mice were on a C57BL/6J background. Because this study incorporated both the *Cthrc1^−/−^* line and lineage-tracing lines, generating fully background-matched littermate controls for every experimental comparison was not feasible. Therefore, we cannot exclude the possibility that differences in genetic backgrounds contributed to some of the observed inflammatory, fibrotic, and calcific phenotypes. Nevertheless, the concordant transcriptomic, histological, and functional findings support the role of *Cthrc1* in AS/CAVD. Future studies using congenic *Cthrc1^−/−^* mice and littermate-controlled cohorts will be required to distinguish the effects of *Cthrc1* deficiency from those attributable to genetic background. Finally, although our findings identified CTHRC1 as a potential mechanistic regulator of valvular remodeling and calcification, we currently lack an effective strategy to deliver CTHRC1 to diseased valvular tissues with sufficient efficiency, specificity, and durability *in vivo*. The development of such delivery approaches, together with their validation in human tissues and complementary experimental models, is an important direction for future investigations.

## Methods

### Mice

The animal experimental protocols were approved by the Institutional Animal Research Committee of Ehime University (approval no. 05RO11-16) and performed in accordance with the institutional guidelines and relevant national regulations for the care and use of laboratory animals in Japan. Seven-week-old male wild-type mice (C57BL/6J background) were obtained from CLEA Japan, Inc., Tokyo, Japan.

Male and female B6D2;129P2-*Cthrc1^tm1Hssk^*/Rbrc (RBRC03519; MGI:3815067) were obtained from the RIKEN BioResource Research Center (Saitama, Japan). We bred these mice to create *Cthrc1^−/−^* mice. All mice were maintained in a mixed C57BL/6 × DBA/2 (B6D2) and 129P2 genetic background. For lineage tracing of *Cdh5*- and *Pdgfrb*-expressing cells, *Tg(Cdh5-cre/ERT2)#Ykub* (59) and *Pdgfrb^tm1.1(cre/ERT2)Csln^* (60) transgenic mice were crossed separately with FVB.Cg-*Gt(ROSA)26Sor^tm1(CAG-lacZ,-EGFP)Glh^*/J (61) reporter mice lacking the LacZ cassette to generate *Cdh5*-CreER^T2^; Rosa26^LSL-GFP^ and *Pdgfrb*-CreER^T2^; Rosa26^LSL-GFP^ offspring, respectively.

All mice were housed in the animal facility of the Graduate School of Medicine, Ehime University, under specific pathogen-free conditions in a 12-h light/12-h dark cycle with controlled temperature (22–24 °C) and humidity (40–60%), and were provided a normal chow diet (MF, Oriental Yeast Co., Ltd., Tokyo, Japan) and water *ad libitum*. The investigators were blinded to allocation during the experiments.

### TTE

To evaluate hemodynamic changes between the groups over time, serial TTE was performed before the procedure and at 3 days, 1 week, 2 weeks, and 16 weeks after the procedure. The mice were examined under 0.5–2.0% isoflurane anesthesia. The mouse chest was shaved to reduce ultrasound attenuation on echocardiography. Mice were gently restrained in the supine position for examination.

Visual Sonic Vevo^®^ 1100 (FUJIFILM, Inc., Tokyo, Japan) was used to measure the AV peak velocity, LV mass, LV volume, and FS. Prior to data collection, the heart rate of the mice was confirmed to be within the range of 580–620 beats per minute. LV dimensions and AV peak velocity were measured using the M and pulse-wave Doppler (PW) modes, with the sample volume tilted at 50°. The turbulence of blood flow through the aortic valve orifice was evaluated using color Doppler mode. The average peak velocity of five consecutive beats was recorded. These measurements were not performed in mice that were judged unable to tolerate restraint or sedation owing to overt postoperative weakness or poor general condition, particularly at 3 days and 1 week after the procedure.

### Establishment of murine aortic valve stenosis model

To establish a murine model of AS/CAVD, we applied an additional wire-mediated injury to the fibrosa-side endothelium of the aortic valve leaflets, based on a previously reported direct WI procedure developed by *Honda et al* (14) (Fig. 1a). Mice were anesthetized with an intraperitoneal injection of a mixture of ketamine (75 mg kg^−1^) and xylazine (10 mg kg^−1^), and their chests were shaved to reduce ultrasound attenuation on echocardiography. Mice were fixed in the supine position, and the right common carotid artery (RCCA) was exposed. A spring guidewire (0.014-inch Miracle 6 PTCA guidewire; AsahiTec Co., Ltd., Aichi, Japan) was introduced into the aortic root via cannulation of the RCCA. The guidewire tip was passed through the aortic valve and maintained in the LV cavity. The ventricular-side surface of the aortic valve was ablated by 30 reciprocal and 50 rotational movements of the spring wire, followed by 50 rotational movements in all Valsalva sinuses to injure the endothelium of both the lamina ventricularis and lamina fibrosa sides of the aortic valves. After the procedure, the spring wire was removed, and the RCCA was ligated to control bleeding. The mice in the sham group underwent simple ligation of the RCCA without contact with the aortic valve leaflets. The skin incision was closed using silk sutures. After the procedure, the mice were placed back into the cage, warmed at 40 °C for 5 h on a heating mat, and then returned to the housing room.

### Isolation of mouse hearts

Mice were euthanized by cervical dislocation, and their hearts were harvested at predetermined time points (6 h, 3 days, 1 week, 2 weeks, and 16 weeks after the WI procedure). Hearts were sectioned along the short axis, immersed in 4% paraformaldehyde, and fixed for 18 h at 22–24 °C. Subsequently, the tissue specimens were processed for paraffin embedding.

### Sample preparation for spatial transcriptome

Four heart specimens per slide were prepared for spatial transcriptomic analysis. The hearts were harvested from C57BL/6J wild-type mice 2 weeks after WI and sham procedure and *Cthrc1^−/−^* mice 2 weeks after WI procedure. Paraffin-embedded mouse heart blocks were prepared as described previously for the isolation of mouse heart sections. An RNA quality check was performed, and the percentage of RNA fragments > 200 nucleotides (DV200) was > 50% in all samples. The blocks were sectioned at a thickness of 10 µm using a microtome and mounted onto the 6.5 mm^2^ oligo-barcoded capture areas on Visium Spatial Gene Expression Slides (10x Genomics, Pleasanton, CA, USA; #PN-2000233), according to the manufacturer’s instructions.

### Visium spatial transcriptomic library preparation and primary data processing

Spatial transcriptomic profiling of FFPE mouse (*Mus musculus*) tissues was performed using the Visium CytAssist Spatial Gene Expression for FFPE assay on a Visium CytAssist instrument (10x Genomics; mouse 6.5 mm kit, #1000445) according to the manufacturer’s protocol (Tissue Preparation Guide, CG000518). FFPE tissue sections were cut at 10 µm, mounted on glass slides, deparaffinized, stained, and imaged according to the manufacturer’s instructions. Probe hybridization and ligation were then performed on the tissue sections, and the resulting ligation products were transferred using the Visium CytAssist instrument to spatially barcoded Visium Gene Expression Slides for library construction according to the manufacturer’s protocol. Libraries were sequenced on a DNBSEQ-G400RS platform (MGI Tech, Shenzhen, China) in the paired-end mode. Primary processing was performed using Space Ranger v2.1.1 (Visium slide/capture-area ID V43M20-207-A1) or v4.0.1 (Visium slide/capture-area ID V45J20-345-A1 and V45J20-345-D1) with default parameters and the transcriptome mm10-2020-A. All delivered downstream analysis files were direct Space Ranger outputs and included cloupe.cloupe, filtered_feature_bc_matrix, filtered_feature_bc_matrix.h5, a spatial folder, and web_summary.html.

Up to four tissue pieces were mounted within each visible-capture area, and three capture areas were processed. Each capture area was initially processed as a single assay, after which the individual tissue pieces were manually realigned in the Loupe Browser to generate section-specific output files for downstream analysis. Tissue pieces with insufficient tissue area on H&E images were excluded to ensure reliable spatial transcriptomic analysis. The final dataset comprised three groups: wild-type sham, wild-type WI, and *Cthrc1^−/−^* WI, with three biologically independent samples per group.

Section-specific outputs were used for subsequent downstream analyses in R using Seurat software.

### Spatial data processing

Raw base call (BCL) files were demultiplexed and aligned using Space Ranger software (v2.1.1 or v4.0.1, 10x Genomics) with the 10x Genomics mouse reference package mm10-2020-A. Image alignment of barcoded spot patterns on the tissue sections and spatial spot selection were performed using the Loupe Browser (v7.0.1, 10x Genomics). The resulting gene-by-spot unique molecular identifier (UMI) count matrices were imported into R (v4.3.3) and processed using Seurat (v5.3.0) (62). Raw UMI counts in the Spatial assay were normalized and variance-stabilized using the SCTransform function (SCTransform v2 with the gamma-Poisson generalized linear model implementation via glmGamPoi, v1.14.3), and the 2,000 most variable genes were identified using the “vst” method. Principal component analysis (PCA) was performed on the normalized expression matrix, and the first 20 principal components were used to construct a shared nearest-neighbor graph and perform graph-based clustering using the Louvain algorithm. Clustering resolutions between 0.1 and 1.0 were explored, and a resolution of 0.4 was chosen for downstream analyses based on cluster stability and biological interpretability. The number of principal components retained was determined using elbow plots, PCA loadings, and dimensionality heatmaps. Low-dimensional embeddings were generated using UMAP, and clusters were visualized in both the embedding space and tissue coordinates using Seurat. To ensure reproducibility, a fixed random seed was used for all analyses involving stochastic steps.

For downstream analyses requiring joint clustering across samples, individual Seurat objects were first generated for each Visium library and annotated with sample-level metadata, including sample identifiers and experimental groups (Seurat metadata columns “orig.ident” and “sample_name”). Cell barcodes were prefixed with the corresponding sample name, and the objects were merged into a single Seurat object. To account for batch and sample-specific effects, we performed integration using the Harmony algorithm (Harmony v1.2.3; R package harmony) implemented via the IntegrateLayers function in Seurat (normalization.method = “SCT,” method = HarmonyIntegration, group.by.vars = “orig.ident”). Harmony embedding was then used as the basis for dimensionality reduction and clustering. UMAP was computed using the first five Harmony dimensions (dims 1–5), and a shared nearest-neighbor graph was constructed using the same dimensions. Louvain clustering was performed again, over a range of resolutions (0.1–1.0), and a resolution of 0.1 was selected for downstream analyses. Unless otherwise specified, analyses requiring integrated representations (e.g., cross-sample visualization and joint cluster definition) were performed on this harmony-integrated object.

### Differential expression analysis

Differential expression analysis was performed at the spot level in R (v4.3.3) using Seurat (v5.3.0) to identify cluster-enriched genes. Individual Visium spots were used as the unit of analysis, and spot identities were assigned according to the Seurat-derived cluster labels (“seurat_clusters”). Pairwise comparisons between clusters of interest were performed using Seurat “FindMarkers” with the default Wilcoxon rank-sum test. Based on histological annotation, cluster 8 corresponded to lesion-associated (stenotic) valve regions, whereas cluster 4 corresponded to intact/normal-like valve regions.

Differential expression testing was performed on the SCT assay using Seurat’s default settings unless otherwise specified. Genes were considered differentially expressed if they showed a Bonferroni-adjusted *p*-value (“p_val_adj”) < 0.05 and an absolute fold change ≥ 1.5. Volcano plots were generated in R using ggplot2 (v3.5.2) by plotting Seurat-reported log2 fold changes (“avg_log2FC”) against −log10 adjusted *p*-values, and selected genes of interest were annotated for visualization.

### Visualization of gene expression

Gene expression patterns were visualized in R (v4.3.3) using Seurat (v5.3.0), ggplot2 (v3.5.2), and ComplexHeatmap (v2.18.0). Violin plots of gene expression were generated using the Vln plot function. The expression of the selected genes in the low-dimensional embedding was visualized using FeaturePlot on the UMAP coordinates, and spatial expression patterns on the tissue sections were visualized using SpatialFeaturePlot. Heatmaps of scaled spot-level expression for selected genes were generated using the ComplexHeatmap package, with columns representing individual visible spots grouped by Seurat cluster.

### Visualization and statistical analysis of module scores

Module scores were calculated at the spot level using the AddModuleScore function in Seurat (v5.3.0). The curated gene sets (Supplementary Data 2 and 3) were matched to the gene symbols detected in the SCT assay and used as the input feature sets. The resulting module scores were visualized across tissue sections using SpatialFeaturePlot. To enable comparison across samples, color scales were harmonized by rescaling scores to the 2nd–98th percentiles of the combined score distribution and applying a common Inferno-color map (viridisLite) with a shared legend. For the statistical comparison of module scores between stenotic and intact valve clusters, analyses were performed at the level of biological replicates. For each module and replicate, we computed the median module score across the spots assigned to clusters 8 (stenotic aortic valve tissue) and 4 (intact aortic valve tissue). Replicates with non-missing median scores for both clusters were retained, and paired differences (cluster 8 minus cluster 4) were evaluated using two-sided Wilcoxon signed-rank tests in R (v4.3.3). Effect sizes were summarized as paired rank-biserial correlations, and 95% confidence intervals (CIs) for the median paired difference were obtained using Wilcoxon tests. For modules tested within the same panel, raw *p*-values were adjusted for multiple testing using the BH FDR procedure.

### GO and GSEA

GO overrepresentation analysis was performed using ClusterProfiler (v4.10.1) in R (v4.3.3). Mouse gene symbols of differentially expressed genes (cluster 8 vs. cluster 4) were converted to Entrez Gene IDs using the bitr function in the org.Mm.eg.db annotation package. GO enrichment was then conducted with the enrichGO function (ontology = “ALL,” keyType = “ENTREZID,” minGSSize = 10, maxGSSize = 500), and results were returned in a readable format (gene symbols). *p*-values were adjusted for multiple testing using the BH procedure, and terms with adjusted *p*-values < 0.05 and *q*-values < 0.2 were considered significantly enriched. The semantic similarity between the enriched GO terms was computed using the pairwise_termsim function, and the top 20 enriched terms were visualized as bar plots, dot plots, and enrichment maps using the barplot, dotplot, and emapplot functions from the enrichplot package. GSEA was performed on a ranked gene list constructed from all genes tested in the differential expression analysis, ordered by log2 FC (avg_log2FC). Mouse GO BP gene sets (MSigDB C5, GO:BP) for *Mus musculus* were obtained using the msigdbr package (v25.1.1), and GSEA was performed with the GSEA function in ClusterProfiler (minGSSize = 10, maxGSSize = 500, p-valueCutoff = 0.05) using target gene sets (full gene lists are provided in Supplementary Data 4 and 5). Enriched pathways were summarized and visualized using dot plots and enrichment maps (emapplots), with color scales indicating either adjusted *p*-values or NES, as implemented in the enrichment plot.

### Statistical analysis

Statistical methods used for spatial transcriptomic analysis and IHC quantification are described in their respective sections. GraphPad Prism (v10.6.1) was used for statistical analyses and graphical representations of all *in vivo* and *in vitro* datasets. TTE and imaging data are presented as mean ± standard deviation or median with interquartile range, unless otherwise specified. Individual data points are shown and represent independent biological replicates, unless otherwise specified.

Data distribution was assessed using the Shapiro–Wilk test. When both groups passed the normality test (*p* < 0.05), an unpaired two-sided Student’s *t* test was performed. If at least one group failed the normality test (*p* < 0.05), the two-sided Mann–Whitney *U* test was used for comparison. For the longitudinal assessment of TTE results, a two-way ANOVA with repeated measures was performed to examine the effects of the intervention group, time, and their interaction. The full model outputs for all the outcomes are summarized in Supplementary Tables 1–12. Imputation was not performed for missing observations owing to scheduled euthanasia or welfare-based omissions at 3 days and 1 week. Post-hoc comparisons were conducted using Sidak’s multiple comparison test. Statistical significance was set at *p* < 0.05.

### Module score calculation and spatial visualization

Gene module activity was quantified for each spot/cell using AddModuleScore in Seurat on the data slot of the SCT assay. Predefined gene sets were matched to the feature names in the Seurat object.

Genes absent from the expression matrix were excluded, and modules with fewer than three matched genes were excluded. Module scores were calculated as the average expression of the input gene set minus the average expression of matched control features using Seurat default parameters, unless otherwise stated (nbin = 24, ctrl = 100, pool = rownames(object), seed = 1). The scores were computed for the combined objects used for each comparison. For between-sample comparisons, spots were assigned to *Cthrc1^−/−^* WI or wild type WI groups according to orig.ident, and biological replicates were defined using sample-level metadata. Spot-level module scores were aggregated to the replicate level by taking the median across spots within each replicate. Replicate-level medians were compared between groups using a two-sided Wilcoxon rank-sum test (exact = FALSE, correct = FALSE). Effect size was summarized as the rank-biserial correlation, and location shift was defined as the difference between the group medians of replicate-level module scores (*Cthrc1^−/−^* WI − wild type WI). Unless otherwise stated, reported *p*-values were unadjusted. Where indicated, multiple-testing correction was performed using the BH FDR method. For spatial visualization, module scores were displayed using SpatialFeaturePlot, with a common color scale across samples and display limits set to the 2nd and 98th percentiles of the score distribution.

### scRNA-seq data processing

Publicly available single-cell RNA-seq datasets were obtained from the NCBI Sequence Read Archive under BioProject accession number PRJNA562645 (SRR10134385–SRR10134390) (25). The data were downloaded as FASTQ files and processed using the CellRanger count pipeline in Cell Ranger (v7.1.0; 10x Genomics) with the prebuilt human reference transcriptome refdata-gen-GRCh38-2020-A (GRCh38; GENCODE v32/Ensembl 98) to generate gene-cell count matrices.

Sample-specific Seurat objects were generated and analyzed in R (v4.3.3) using Seurat (v5.3.0). Genes detected in fewer than three cells were excluded from analysis. Cells with 200 or fewer detected genes, 4,000 or more detected genes, or a mitochondrial transcript proportion of 5% or greater were excluded. Subsequently, six filtered sample-level objects were merged.

The UMI counts were normalized and variance-stabilized using SCTransform with v2 regularization and glmGamPoi-based model fitting. PCA was performed using the variable features identified by SCTransform. The processed Seurat data were subsequently converted into a Monocle 3 object for dimensionality reduction, clustering, subclustering, and trajectory analysis.

### Single-cell clustering and cell-type annotation

Following preprocessing in Seurat, raw UMI counts and the corresponding cell- and gene-level metadata were used to construct a Monocle 3 (v1.4.26) “cell_data_set” object. PCA was performed in Monocle 3 using 3,000 variable genes identified by SCTransform, and the first 50 principal components were retained. Sample-associated technical variation was mitigated using mutual nearest-neighbor alignment implemented in “align_cds(),” with sample identity specified as the alignment group. A UMAP embedding was generated from the aligned coordinates, and cells were clustered using the Leiden algorithm at a resolution of 1 × 10^−3^. Cell type identities were assigned based on the expression of established marker genes and visualized using feature and dot plots.

### Trajectory and pseudotime analysis

For focused trajectory analysis, the cells assigned to Monocle 3 partition 1 and selected preliminary Seurat clusters were retained. The resulting Monocle 3 cell_data_set was reclustered at a resolution of 5 × 10^−4^, and a principal graph was learned using learn_graph(). Root principal nodes were manually selected within the *TGFBI*-high, *COL1A1*-high, and *COL3A1*-high cell populations (Fig. 4i, j and Supplementary Fig. 7a, b), which were designated as the starting state for pseudotime ordering. The cells were then ordered along the trajectory using order_cells(). Pseudotime was visualized on the Monocle 3 UMAP embedding.

Pseudotime-resolved expression patterns of selected genes were descriptively visualized by plotting gene expression values against Monocle 3-derived pseudotime using “plot_genes_in_pseudotime().” The fitted expression trends were overlaid using the default natural spline function of pseudotime, and no statistical testing was performed for these descriptive plots.

### Human tissue specimens and IHC

Archival de-identified human aortic valve specimens were obtained from the same specimen set previously reported by Komoda *et al* (63). Pathological and non-diseased control specimens were obtained from patients undergoing SAVR at the Ehime University Hospital. DAB IHC was performed on pathological (n = 7; surgically resected specimens) and non-diseased control specimens (n = 7; autopsy specimens).

The use of these human specimens was approved by the Ehime University Internal Review Board (Protocol Nos. 1509022 and 1603002). Detailed patient and specimen characteristics for this specimen set have been reported in the Supplementary Information section of the original report (63). DAB^+^ staining was quantified as described in the immunostaining section. The same threshold criteria were applied to all the samples. The percentage of DAB^+^ areas was compared between the pathological and non-diseased control specimens using a two-sided Mann–Whitney *U* test.

### Immunostaining

Murine hearts were fixed in 4% paraformaldehyde phosphate buffer solution for 18 h at room temperature, paraffin-embedded, and sectioned at 3-μm thickness. Sections were deparaffinized and rehydrated, and heat-induced antigen retrieval was performed in 0.01 M citrate buffer (pH 6.0) using an autoclave at 125 °C for 15 min before immunostaining.

For immunofluorescence, after antigen retrieval, the sections were incubated with the primary antibodies listed in Supplementary Data 6, diluted in Dako REAL^TM^ Antibody Diluent (Agilent Technologies, Singapore; # S2022), for 2 h at room temperature, without an additional blocking step. Sections were then washed three times in PBS for 3 min each, incubated with the appropriate secondary antibodies listed in Supplementary Data 6 for 2 h at room temperature, washed under the same conditions, counterstained with 0.5 µg/mL Hoechst 33342 for 30 min, washed again with PBS as described above, and mounted with ProLong^TM^ Gold Antifade Mountant (Invitrogen, Thermo Fisher Scientific #P36930). Fluorescent images were acquired using an A1Rsi-E1 confocal microscope (Nikon Corporation, Tokyo, Japan).

Following antigen retrieval, endogenous peroxidase activity was quenched in 3% H2O2 for 15 min. Sections were washed thrice in PBS for 3 min each, incubated with the primary antibodies listed in Supplementary Data 6 for 2 h at room temperature, washed again under the same conditions, and then incubated with the appropriate horseradish peroxidase-conjugated secondary antibodies provided in Supplementary Data 6 for 45 min at room temperature. Immunoreactivity was visualized using a Histofine DAB Substrate Kit (catalogue no. 425011; Nichirei Biosciences, Tokyo, Japan) as detailed in Supplementary Data 6. Bright-field images of the DAB-stained sections were acquired using a BZ-X800 microscope with the BZ-X Viewer software (version 1.3.0.1; KEYENCE, Osaka, Japan). The DAB^+^ area fraction was quantified using BZ-X Analyzer software (version 1.1.30.19; KEYENCE, Osaka, Japan), with the total area of the aortic valve used as the reference area. Thresholds were set separately for each antibody and applied consistently across all corresponding samples. The DAB^+^ area was normalized to the total aortic valve area.

## Abbreviations used

ADRES: Advanced Research Support
ANOVA: analysis of variance
AS: aortic stenosis
BP: Biological Process
CAVD: calcific aortic valve disease
CI: confidence intervals
DGE: differential gene expression
EndMT: endothelial-to-mesenchymal transition
FDR: false discovery rate
FS: fractional shortening
GEO: Gene Expression Omnibus
GO: Gene Ontology
GSEA: gene set enrichment analysis
H&E: Hematoxylin & Eosin
IPA: Ingenuity Pathway Analysis
LV: left ventricular
NES: normalized enrichment score
PCA: principal component analysis
PW: pulse-wave
RBC: red blood cells
RCCA: right common carotid artery
RIMD: Research Institute for Microbial Diseases
SAVR: surgical aortic valve replacement
TAVI: transcatheter aortic valve implantation
UMAP: uniform manifold approximation and projection
UMI: unique molecular identifier
VEC: valvular endothelial cells
VIC: valvular interstitial cells
WI: wire injury

## Reporting summary

Further information on the research design is available in the Nature Portfolio Reporting Summary linked to this article.

## Data availability

The raw and processed Visium spatial transcriptomic data generated in this study have been deposited in the NCBI Gene Expression Omnibus (GEO) under the accession code GSE336239. This dataset will be made publicly available upon publication. All other data supporting the findings of this study are available from the corresponding author upon request.

## Code availability

No new software or algorithms were developed for this study. All computational analyses were performed using publicly available software packages as described in the Methods section. The R scripts used for data analysis and figure generation are available from the corresponding author upon request.

## Acknowledgements

We acknowledge Daisuke Motooka from the Research Institute for Microbial Diseases (RIMD), Osaka University, Osaka, Japan and Mei Miyazaki from the Advanced Research Support Center (ADRES), Ehime University, Ehime, Japan, for their technical support. Additionally, we would like to thank Editage (www.editage.jp) for English language editing.

## AUTHOR CONTRIBUTIONS

Conceptualization: T.S. and H.S.; Methodology: T.S., H. M., H.S., and D.S.; Investigation: H.S., T.S., and M.K.; Formal analysis: H.S., T.S., and D.S.; Data curation: H.S. and T.S.; Software: D.S. and H.S.; Validation: H.S. and T.S.; Visualization: H.S. and T.S.; Resources: T.S. and Y. K. (Yoshiaki). Funding acquisition: T.S., H.S., H.I.; Project administration: H.S., T.S., H.I.; Supervision: T.S., H.I.; Writing–original draft: H.S.; Writing–review and editing: T.S., M.H., Y.N., M.K., A.I., H.K., Y.K. (Yosuke), F.T., H.K., T.N., S.U., D.S., Y.K. (Yoshiaki), O.Y. and H.I.

## Novelty and significance

### What Is Known?

* Calcific aortic valve disease (CAVD) is a progressive disease in which inflammation, tissue remodeling, and mineral accumulation gradually disrupt aortic valve function, but the endogenous mechanisms that protect the valve tissue after endothelial injury and restrain subsequent calcification remain poorly understood.

* Valvular interstitial cells (VICs) are generally viewed as important regulators of valve remodeling, but their potential role in mounting tissue-protective responses during early disease has received less attention.

* Macrophages accumulate in diseased aortic valves and are associated with inflammatory remodeling, but how the resident valvular stromal cells influence this inflammatory environment during the transition from valve injury to calcification remains unclear.

### What New Information Does This Article Contribute?

* This study identifies CTHRC1 as a prominent component of an early injury-induced stromal response, and detects its expression in expanded VICs derived predominantly from PDGFRβ^+^ resident interstitial cells and also in human stenotic aortic valves.

* *Cthrc1* loss exacerbates chronic valvular calcification and is accompanied by accumulation of lipid-laden macrophages and evidence of greater cellular stress and injury within the valve.

* These findings reveal that injury-activated VICs can function as a tissue-defense compartment, helping maintain the valve microenvironment and limit inflammatory remodeling that precedes calcification.

## Summary Narrative

CAVD is characterized by progressive inflammatory and fibrocalcific remodeling, but the endogenous mechanisms that protect the valve from progression after initial tissue injury are poorly understood. This study identifies an injury-induced, CTHRC1-dependent stromal response that protects the valvular interstitial microenvironment during the early response to injury. After endothelial injury, CTHRC1 is transiently induced in expanded VICs derived predominantly from resident PDGFRβ^+^ interstitial cells and its loss is associated with pronounced accumulation of foamy macrophages, impaired mitochondrial respiratory-chain signatures, and activation of cell death– associated pathways. Importantly, this protective stromal response is identified in an injury-driven model without experimentally induced dyslipidemia and is supported by CTHRC1 expression in human stenotic aortic valves. These findings extend the current understanding of VICs beyond their established roles in pathological remodeling and identify the valvular stroma as an active tissue-defense compartment that restrains inflammatory injury and dystrophic calcification. More broadly, this framework provides evidence that endogenous stromal responses can regulate the inflammatory microenvironment after tissue injury and influence CAVD progression, suggesting CTHRC1-dependent stromal protection as a potential therapeutic axis for limiting disease progression.

